# ADAMTS3 as a promising novel biomarker for the diagnosis of hepatocellular carcinoma

**DOI:** 10.1101/2025.09.17.676767

**Authors:** Chen Wang, Huaizhou Jin, Qijie Zhang, Yanqiu Zou, Yaosen Wang, Yanlong Cai, Zhuo Fang, Li Yao, Nicolò Maccaferri, Ali Douaki, Jianguo Feng, Denis Garoli, Shangzhong Jin

## Abstract

Early diagnosis of hepatocellular carcinoma (HCC) still represents a significant challenge. The rising of obesity and non-viral liver disease associated HCC further limits the effectiveness of conventional screening approaches such as ultrasonography and alpha-fetoprotein (AFP) testing. In this context, the identification of potential effective biomarkers for rapid and precise diagnosis of HCC still represents an extremely important task. Here, we identify ADAMTS3 as a novel mechanistic serum biomarker for HCC that addresses this shifting etiological landscape. Transcriptome analysis of four independent cohorts–including HCC patients, individuals with obesity-or non-viral liver disease, and healthy controls– reveals four secreted proteins associated with HCC progression. Among them, ADAMTS3 demonstrates strong diagnostic performance by ROC (receiver operating characteristic) and LASSO (least absolute shrinkage and selection operator) analysis. High ADAMTS3 expression correlates with aggressive molecular signature, characterized by enhanced proliferative signaling, suppressed immune effector responses, and a reprogrammed immune microenvironment marked by reduced NK cell infiltration and increased accumulation of immunosuppressive tumor-associated macrophages, driven by extracellular matrix stiffening. Knockdown of ADAMTS3 in cell lines significantly reduced proliferation, and clonogenic potential. Reintroduction of ADAMTS3 partially rescued both phenotypes, further confirming its role in promoting HCC cell growth. Optical tweezer–based measurements further reveal reduced cell stiffness upon ADAMTS3 deficiency, highlighting its role in extracellular matrix (ECM) remodeling. Finally, to provide a simple strategy for HCC diagnosis based on this novel biomarker, we demonstrated a DNA origami-based SERS biosensor capable of detecting ADAMTS3 at 10^-11^ M. Together, these findings identify ADAMTS3 as a mechanistic biomarker and demonstrate a translational sensing strategy for non-invasive HCC early screening.

## 1. Introduction

Liver cancer ranks as the third leading cause of cancer-related deaths worldwide, with hepatocellular carcinoma (HCC) representing about 80% of cases^1^. Unlike most other major cancers, mortality rates for HCC have shown improvement over the past decade^2^. For patients with early-stage disease (Barcelona Clinic Liver Cancer stage A), surgical options remain the treatment of choice, offering curative potential: median overall survival (OS) exceeds 5 years following resection and can surpass 10 years after liver transplantation^3,4^. Unfortunately, nearly 70% of patients are diagnosed at an advanced stage, when curative therapies are no longer viable^2,5^. This highlights the urgent global need for more effective strategies enabling earlier detection of HCC.

Abdominal ultrasonography is the primary screening tool currently recommended by major international liver associations^6^. However, its diagnostic performance is highly dependent on operator expertise and is notably less effective in detecting small HCC nodules or in patients with obesity or non-viral liver disease^7–11^. To mitigate these limitations, serum biomarker testing is often used in combination with ultrasonography. Biomarker-based screening reduces operator dependence, improves sensitivity, and offers greater convenience— factors that can strengthen diagnostic accuracy, promote patient adherence, and facilitate long-term surveillance. Among available biomarkers, alpha-fetoprotein (AFP) remains the most widely used in clinical practice for HCC detection. Nonetheless, AFP has important limitations: its levels may remain normal even in patients with large tumors and can also be elevated in benign liver conditions^12^. demonstrated potential in specific clinical settings. For example, des-gamma carboxyprothrombin (DCP) has shown strong diagnostic performance in distinguishing HCC from hepatitis B virus–related chronic liver disease (sensitivity: 84.08%; specificity: 90.43%)^13^. Similarly, ST6GAL1 has been identified as a promising biomarker for predicting susceptibility to Lenvatinib therapy^14^. Despite these advances, no universally reliable serum biomarker for early HCC detection has yet been established, particularly in patients with liver nodularity associated with obesity or non-viral etiologies. This gap is of growing importance given the epidemiological transition in HCC burden—from virus-related disease to causes such as alcohol-associated liver injury and obesity.^12^.

To advance early detection and improve management of HCC in broader populations, we performed a transcriptome-based screening across four independent cohorts—comprising healthy controls, individuals with obesity or non-viral liver disease, and HCC patients with diverse etiologies—to identify serum-secreted proteins linked to HCC progression. This analysis revealed four consistently upregulated candidates: ADAMTS3, MAPT, CFB, and PRH2. Among them, ADAMTS3 emerged as the most promising biomarker, demonstrating strong discriminatory capacity in both early-stage and AFP-negative HCC. Notably, ADAMTS3 also acts as a mechanistic biomarker that actively promotes HCC progression. Although relatively underexplored, ADAMTS3 is localized at or near the cell surface, where it functions as the principal N-propeptidase of procollagen II—initiating collagen fibril assembly, the key structural component of the extracellular matrix (ECM)—and as a VEGF-C–activating protease, thereby regulating lymphangiogenesis and angiogenesis. ^15,16^. In cancer biology, limited studies suggest that ADAMTS3 plays diverse roles, acting as a tumor promoter in glioma stem cells but exhibiting tumor-suppressive effects in breast cancer. Its function in HCC, however, remains largely unclear and has not been systematically explored. In this study, we demonstrate that ADAMTS3 remodels the extracellular matrix, fostering an immunosuppressive microenvironment and activating GPCR signaling pathways that enhance stemness and tumorigenic potential.

Conversely, serum biomarkers are usually present at very low concentrations, typically within the picogram to nanogram per milliliter range^17,18^. Therefore, the ability to accurately detect ADAMTS3 against the complex serum background is crucial. At present, serum biomarker detection primarily relies on immunoassays such as Enzyme-Linked Immunosorbent Assay (ELISA) and Chemiluminescence Immunoassay (CLIA).^19,20^. However, ELISA is limited by suboptimal sensitivity, a narrow dynamic range, and vulnerability to background interference. While CLIA provides improved sensitivity and greater automation compared to ELISA, it remains reliant on expensive instrumentation and proprietary reagents, and still suffers from nonspecific signal noise. In contrast, Surface-Enhanced Raman Scattering (SERS) represents a powerful alternative, offering ultrahigh sensitivity—reaching femtomolar or even single-molecule levels—along with molecular fingerprint-level specificity. Owing to these features, SERS has been widely explored for biomarker-based early cancer diagnosis, benefiting from its exceptional sensitivity and noninvasive nature. However, its practical application is frequently limited by the need for stable electromagnetic “hotspots,” which are the dominant contributors to SERS signal enhancement—surpassing chemical amplification by several orders of magnitude—and are critical for ensuring reliable detection. ^21^.

To address this limitation, stable and robust SERS “hotspots” can be generated through precisely controlled nanoparticle assembly methods. Among these, DNA origami technology— first introduced by Paul Rothemund in 2006—has emerged as a versatile platform for the programmable folding of DNA into defined shapes and patterns. This enables the construction of nanoscale architectures with precise spatial control, allowing the organized arrangement of plasmonic nanoparticles to produce reproducible plasmonic “hotspots.” In addition, DNA origami structures are highly adaptable to chemical modification, making them particularly suitable for analytical applications.^22^.

In this study, recognizing the shifting landscape of HCC etiology, we identified ADAMTS3 as a novel mechanistic serum biomarker by leveraging high-throughput RNA sequencing data from healthy individuals and patients with obesity, non-alcoholic fatty liver disease (NAFLD), virus-associated HCC, and both early-and late-stage HCC. Building on this discovery, we further developed a DNA origami–based immunosensor capable of ultrasensitive and highly specific detection of ADAMTS3, offering clinical promise for noninvasive testing from just a single drop of blood.

## 2. Materials and Methods

### 2.1 Differential expression analysis

The data used in this study are publicly available (GSE164359, GSE141200, GSE126848, GSE199819, PRJNA912860, PRJNA715360, PRJNA743855). All corresponding analyses were performed using R (version 4.4.1). The Differential expressed genes (DEGs) were identified through the application of the DESeq2 package, with the cutoff criteria (adjusted p-value < 0.05 and |log_2_FoldChange|> 1). The diagnostic performance of candidate genes was assessed using the pROC package. Candidates subjected to LASSO (least absolute shrinkage and selection operator) analysis, and the LASSO regression model were implemented with the glmnet package. Identified DEGs were subsequently analysed for Gene Ontology (GO) terms and Kyoto Encyclopedia of Genes and Genomes (KEGG) pathway enrichment using the clusterProfiler package, and the results were visualized with ggplot2 package. A significance threshold of p < 0.05 was applied throughout the analyses.

### 2.2 Cell lines

The Huh-7, and HEK293T cells were obtained from American Type Culture Collection. HEK293T and Huh-7 were maintained in DMEM media with 10% FBS and P/S. All cells were maintained at 37 °C in a humidified atmosphere with 5 % CO2 and all the experiments were performed with cell lines cultured for < 4 months.

### 2.3 Preparation of expression vectors

3xFlag-ADAMTS3 expression plasmids were generated using a Hieff Clone® Universal II One Step Cloning Kit (Yeasen Biotechnology, China), according to the manufacturer’s protocol. Primers used to construct Flag-tagged ADAMTS3 are listed below.

Fwd-flag: accggcgcctactctagagctagcgaattcgccaccatggactacaaagacc; Rev-flag: aaccaaagtgacaggagaaccatggatcccttgtcatcgtcatccttgtaa; Fwd-ADA: atggttctcctgtcactttggttg, Rev-ADA: ggggagggagaggggcgcggccgcttatctttctaaggtggatgatc.

Backbone vectors were digested with NheI and NotI restriction endonucleases (New England Biolabs). E. coli DH5a (Yeasen Biotechnology, China) was used for plasmid cloning. shRNAs targeted for ADAMTS3 were cloned into AgeI and EcoRI digested piggybac-U6 vector with NEB Quick Ligation™ Kit. Corresponding sequence for shRNAs are listed in Table S1.

### 2.4 Proliferation Assay

Cell Counting Kit-8 (CCK-8, APExBIO) was used to perform proliferation assays. In brief, 1.5×10^3^ cells were plated in 100µL normal growth medium in 96-well plates and 10 µL of the CCK-8 solution was added to each well. The absorbance at 450 nm was measured using a microplate reader (Multiskan FC) after incubation for 1 h.

### 2.5 Colony Formation Assay

800 cells per group were seeded in six-well plates in 2 ml of medium. Fresh medium was added after 96 hours to prevent nutrient exhaustion. At endpoint, colonies were fixed with 4% Paraformaldehyde Fix Solution (Beyotime, P0099) for 10 min at room temperature, then washed twice with PBS. The samples were subsequently stained with crystal violet solution (Beyotime, C0121) at 37 °C for 10 minutes and rinsed thoroughly with water. ImageJ was used for quantification of area covered by colonies and colony number.

### 2.6 Western Blot Analysis

5 x 10^5^ cells/well were seeded into six-well plates and incubated for 48 h. Cell samples were rinsed thoroughly with PBS. Total cellular protein was extracted on ice using Mammalian Protein Extraction Reagent (M-PER™, Thermo Scientific) supplemented with Halt™ Protease Inhibitor Cocktail. Supernatant were then collected by centrifugation at 18,000 × g for 10 min at 4 °C. Cell extracts were diluted with loading buffer to a final protein concentration of 1 mg/mL, separated by SDS-PAGE, and transferred onto polyvinylidene difluoride (PVDF) membranes (Bio-Rad). The membranes were then blocked with 5% non-fat milk. After 3 washes with TBST, membranes were reacted with target antibody overnight at 4 °C. The membranes were further washed with TBST and incubated with a horseradish peroxidase– conjugated secondary antibody for 1 h at room temperature. After another wash step, the membranes were reacted with BeyoECL Plus working solution (Beyotime, P0018S).

### 2.7 DNA Origami Assembly

DNA staple strands (0.1 µM each) were pooled by combining 0.5 µL from each tube into a total volume of 101 µL (sequences are listed in Supplementary Table S1). The pooled mixture was vortexed thoroughly and diluted to 1% by mixing 1 µL of the pooled staples with 99 µL of ultrapure water. To assemble DNA origami, an annealing solution (100 µL total) was prepared by combining 20 µL of the diluted staple mixture, 10 µL of 10× TAE buffer supplemented with 0.1 M MgCl_2_, 67.5 µL of ultrapure water, and 2.5 µL of m13mp18 scaffold (250 µg/mL). The solution was subjected to a thermal annealing protocol, which consisted of heating to 80°C followed by gradual cooling to 20 °C. Assembled DNA origami structures were purified by agarose gel electrophoresis (1% agarose in 1× TAE containing 0.1 M MgCl_2_) at 75 V for 60 min. The desired DNA band was excised and extracted using the SanPrep Column DNA Gel Extraction Kit, yielding 25 µL of purified DNA origami.

### 2.8 DNA Origami-Gold Nanoparticle Conjugation

25 µL of purified DNA origami was incubated with 2 µL of TCEP to reduce and activate thiol groups. In parallel, 100 µL of gold nanoparticle solution was centrifuged at 2900 × g for 5 min. The supernatant was discarded, and the pellet was resuspended in 26.5 µL of ultrapure water containing 3.5 µL of 0.2% SDS. The activated DNA origami was then mixed with the prepared gold nanoparticles and incubated at 40°C for 40 min without agitation. To facilitate salt-aging, NaBr was sequentially added at 10-min intervals with gentle shaking in the following volumes and concentrations: 1.7 µL (0.4 M), 2.1 µL (0.4 M), 2.3 µL (0.4 M), 2.3 µL (1 M), 2.3 µL (1 M), 2.3 µL (1 M), and 6.5 µL (1 M). This was followed by the addition of 6 µL and 6.6 µL of 1× TAE buffer containing 5 mM MgCl_2_ at 10-min intervals. The conjugated product was purified through four washing steps, each involving centrifugation at 2900 × g for 5 min, removal of the supernatant, and resuspension in 20 µL of fresh buffer. The first two washes were performed using 1× TAE with 5 mM MgCl_2_ and 0.02% SDS, while the final two washes used 1× TAE with 5 mM MgCl_2_ without SDS.

### 2.9 Raman spectroscopy Detection

DNA origami-gold nanoparticles were incubated with either 1X PBS (blank) or purified Histagged-ADAMTS3 at diverse concentrations. A 10 µL aliquot of each mixture was deposited onto a silicon wafer and subjected to Raman spectroscopy. Measurements were performed on a Horiba XploRA Plus Raman microscope equipped with a 50× long-focal-length objective (NA = 0.75). Visible Raman spectra were recorded using a 633 nm excitation laser set at 3.5% power.

### 2.10 Measurement Young’s modulus by optical tweezer

The mechanical properties of cells were measured using the Lumicks C-Trap optical tweezer system^23,24^. Polystyrene beads (∼2 μm in diameter; 2.5 wt%) were diluted 1:1000 in 1X PBS and added to a chamber (1 mm × 1 cm²) containing a single layer of adherent cells. Individual beads were trapped and displaced using a calibrated laser, while the corresponding force and displacement were recorded in real time. Force–indentation curves were generated and analyzed using the standard Hertz model to calculate the Young’s modulus of each cell. Data acquisition and analysis were performed using the manufacturer-provided software and custom MATLAB scripts.

### 2.11 Synthesis of AuNPs

AuNPs were synthesized using a modified Turkevich–Frens method, as detailed by Smirnov^25^. In brief, an aqueous solution of hydrogen tetrachloroaurate (III) was heated to boiling under continuous stirring, and a reducing agent—trisodium citrate, potassium ascorbate, or ascorbic acid—was added dropwise. This reduction led to the formation of colloidal gold, with particle size and distribution controlled by the type and concentration of the reducing agent. The resulting nanoparticles were stable in colloidal suspension and suitable for subsequent biosenor construction.

### 2.12 Numerical simulations

Numerical simulations were carried out to investigate the electromagnetic field distribution close to the dimer nanoparticle system. To this aim, the electromagnetic response of an isolated (non-interacting) dimer in solution was simulated using the Finite-Element Method (FEM) implemented in the Wave Optics Module of Comsol Multiphysics®. The radius of the single building block of the dimer (sphere) was set according to the average nanoparticles’ sizes obtained during SEM investigations (50 nm), while the interparticle gap was set to 2.5 nm. To consider the presence of the solvent, the refractive index of the environment was set to be n=1.33 (water). The unit cell was set to be 200 nm wide in both x-and y-direction (dimer plane, with dimer axis aligned along the y-direction) and 2000 nm in z-direction, with perfect matching layers (500 nm thick) at the borders. The incident light is an unpolarized plane wave propagating along the z-direction.

### Key resources table

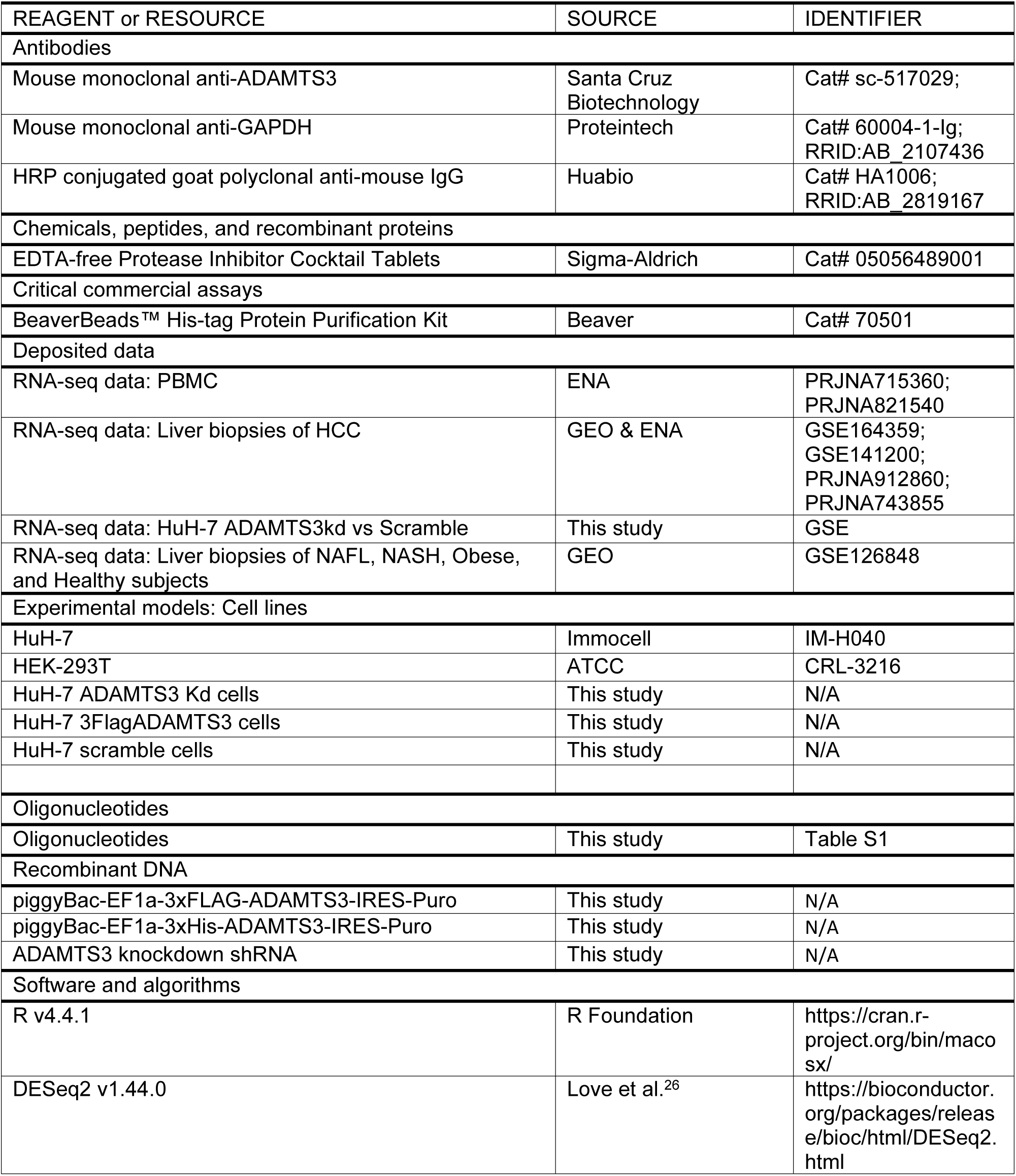
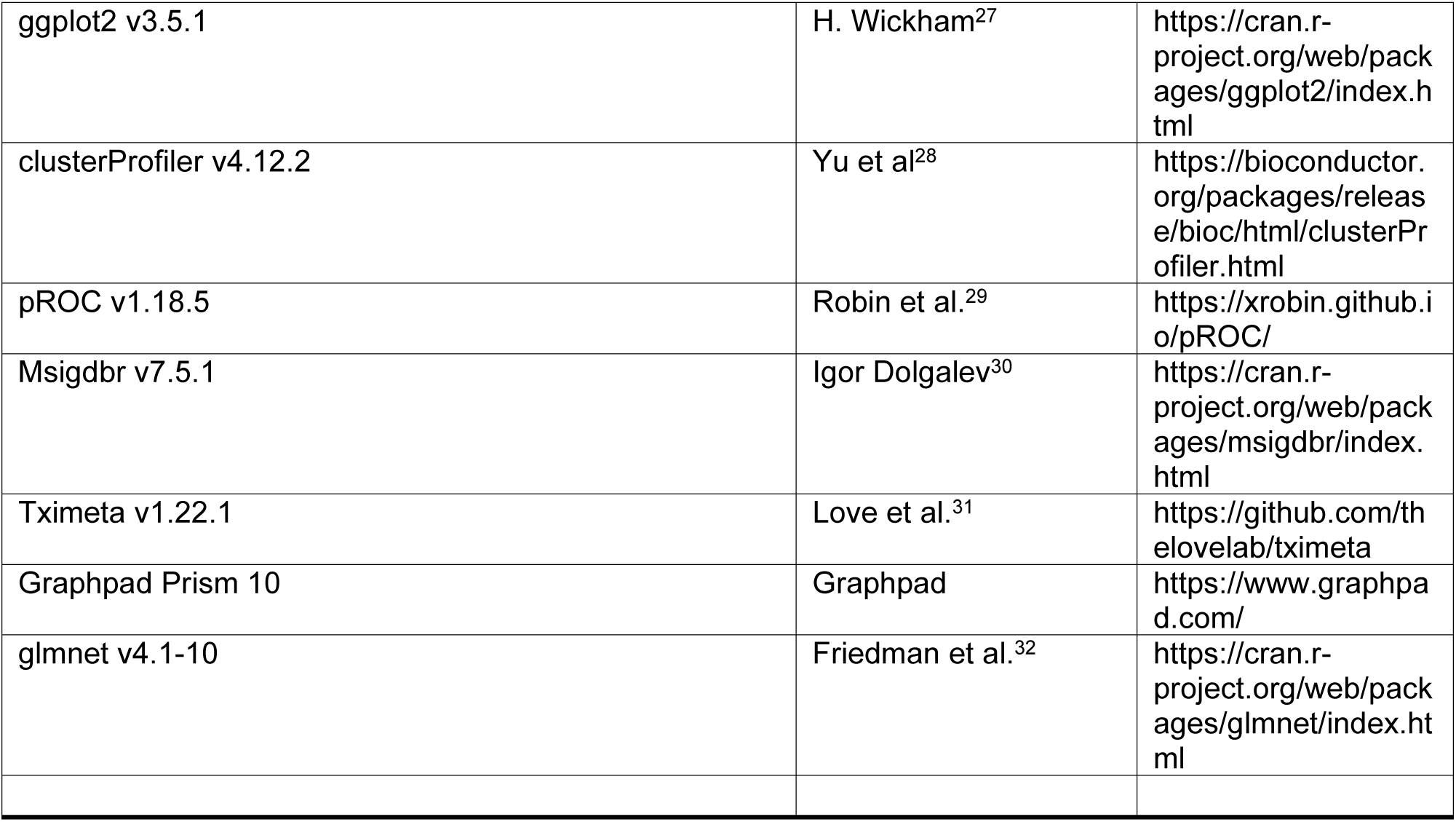

## 3. Results and Discussion

### 3.1 Identification and Validation of ADAMTS3 as a Serum Biomarker in HCC Patients

To identify a broadly applicable and specific serum biomarker for HCC patients, we first analyzed molecular profiles from a discovery cohort comprising non-HCC individuals— including healthy controls and patients with obesity or NAFLD—as well as HCC patients with diverse etiologies (Hepatitis B virus, Hepatitis C virus, and non-viral) and disease stages (Figure 1).

**Fig. 1.**
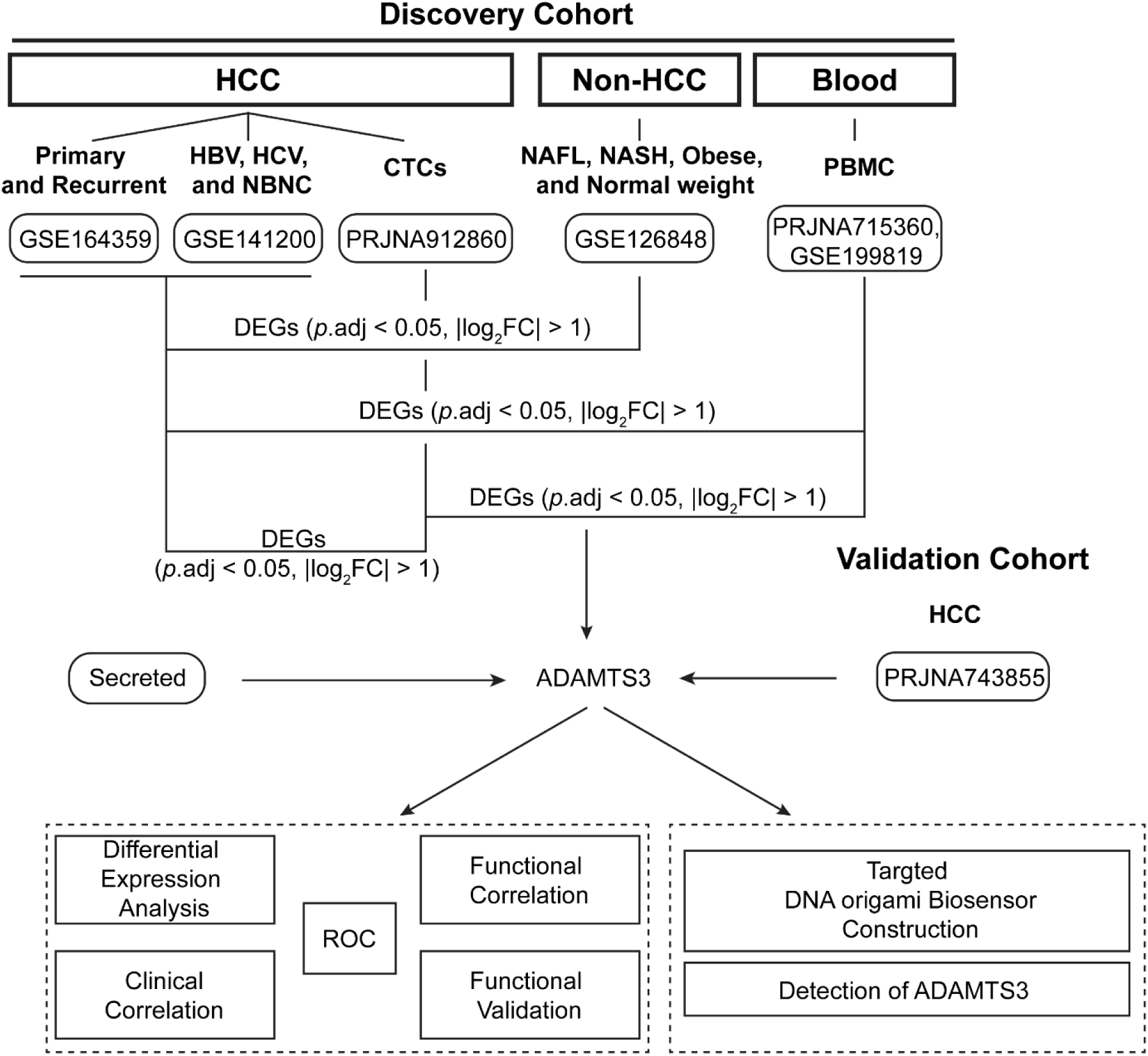
The flowchart of data collection, analysis, and experimental validation.

By performing differential expression analysis comparing liver tissues of HCC to non-HCC (*p*.adj < 0.05, |Log_2_FoldChange| > 1), 7665 genes were observed as significantly up-regulated and 3719 genes were exhibited significantly down-regulation (Figure 2A). Another RNA profile comparison between HCC tissues and peripheral blood mononuclear cells (PBMCs) (*p*.adj < 0.05, |Log_2_FoldChange| > 1) identified 5,681 up-regulated and 3,033 down-regulated genes (Figure 2B). Additionally, differential analysis comparing circulating tumor cells (CTCs) to HCC tissues identified 1,861 upregulated and 287 downregulated genes (Supporting Information - Figure S1A). All upregulated genes were further filtered using a threshold of |Log_2_FoldChange| > 1.5, intersected across comparisons, and then cross-referenced with secreted proteins annotated in the GeneCards database. This yielded four candidate genes that are both secreted and consistently up-regulated across the discovery cohort (Figure 2C). As these candidates represent promising serum biomarkers for HCC detection and monitoring, we further evaluated their performance in distinguishing HCC from health individuals with ROC (receiver operating characteristic) analysis. Among these candidates, ADAMTS3 and MAPT showed the highest area under the curve (AUC) values in ROC analysis, indicating superior diagnostic performance (Figure 2D). For diagnosis biomarkers, achieving an optimal balance between specificity (i.e., true negative rate) and sensitivity (i.e., true positive rate) remains a significant challenge ^33^. In clinical practice, tests with high specificity are typically prioritized to reduce false positives. Accordingly, we focused on the partial AUC within the 90–100% specificity range for ADAMTS3 and MAPT ^29^. However, no significant difference was observed between the two markers in this high-specificity range (*p* = 0.2105). Further analyses using the LASSO regression algorithm revealed that ADAMTS3, rather than MAPT, exhibited highest predictive value (Figure 2E & F). Therefore, we focused on ADAMTS3 for further investigation. We assessed the predictive value of ADAMTS3 in datasets comprising primary HCC versus healthy controls, as well as recurrent HCC versus healthy controls. Notably, ADAMTS3 maintained reliable predictive performance, achieving an AUC of 0.9039 (95% CI: 81.79%–99%) for primary HCC and 0.8704 (95% CI: 77.32%–96.75%) for recurrent HCC (Figure 2G). We also evaluated the predictive value of ADAMTS3 in more challenging clinical contexts. Specifically, we assessed the performance of ADAMTS3 in obesity/NAFL and HCC populations. ADAMTS3 consistently demonstrated strong discriminatory power, with an AUC of 0.9062 (95% CI: 83.67%–95.60%) in distinguishing HCC from obesity and NAFL cases (Supporting Information - Figure S1B). Furthermore, to better address the limitation of AFP in identifying patients with low AFP expression^34^, we tested the predictive performance of ADAMTS3 in HCC patients within the bottom quartile of AFP levels. ADAMTS3 remained a robust predictor in this subgroup, showing consistent performance, with an AUC of 0.8765 (95% CI: 78.36%–96.94%) (Figure 2H). To validate these findings, we next assessed ADAMTS3 performance in an independent dataset. Notably, ADAMTS3 showed strong predictive ability also in this cohort, with an AUC of 0.9038 (95% CI: 84.69%-96.06%) (Figure 2I). Further, we analyzed ADAMTS3 expression across different stages of HCC progression. No significant difference was observed between the obesity/NAFL and healthy groups, whereas ADAMTS3 levels were significantly elevated in HCC compared to both healthy and obesity/NAFL groups (Figure 2J). ADAMTS3 expression among PBMCs, normal liver tissues, obese/NAFL liver tissues, HCC tissues, and metastatic samples (CTCs) progressively increased with disease progression (Figure 2J). In addition, mapping molecular features to biological behaviors, we observed a significant enrichment of M2 macrophages and a corresponding exclusion of Natural killer (NK) cells in the tumor microenvironment of the high ADAMTS3 group (Figure 2K). Finally, 10-year overall survival was significantly reduced in patients with high ADAMTS3 expression (Figure 2L).

**Fig. 2.**
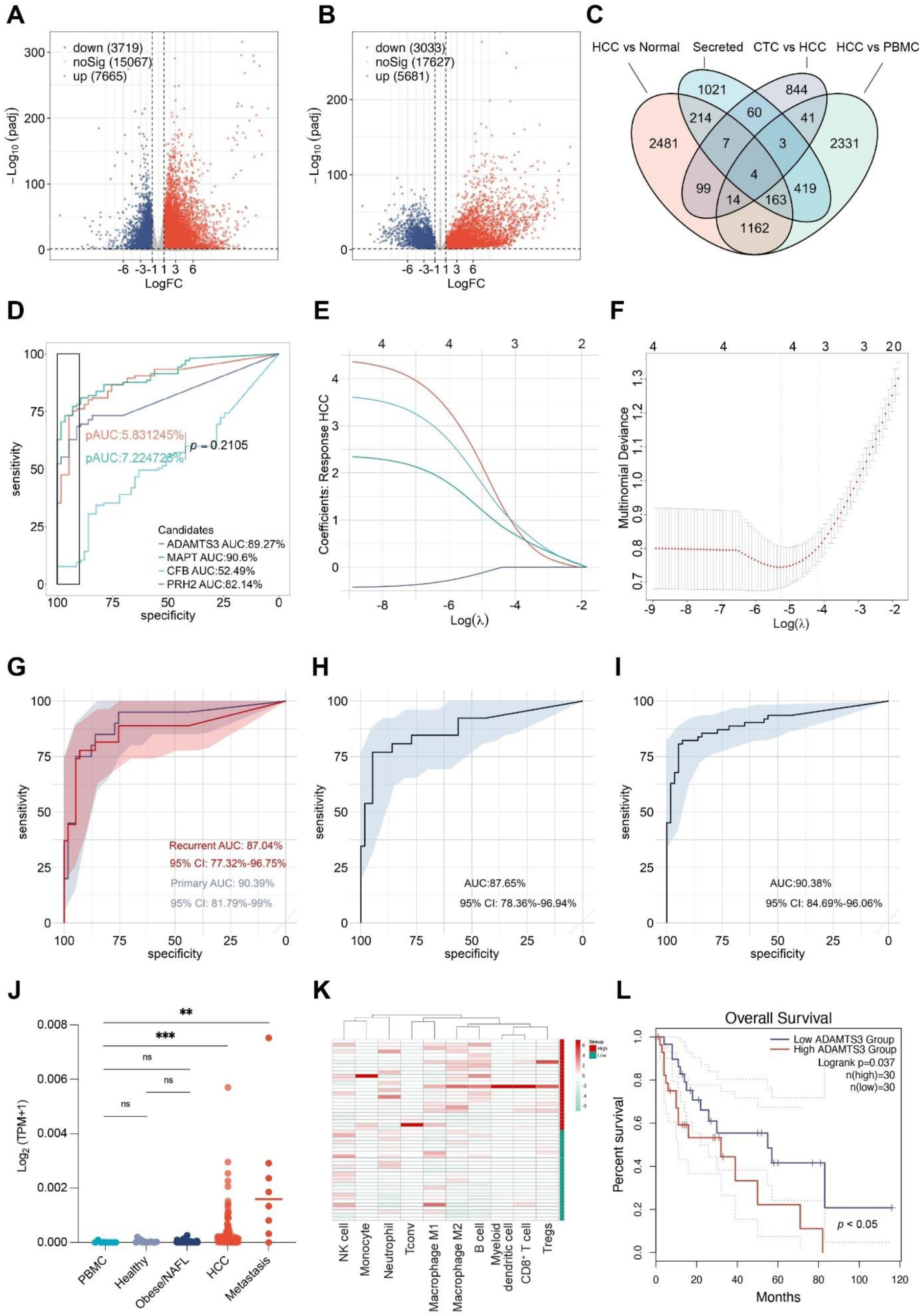
Identification and Validation of ADAMTS3 as a Serum Biomarker in HCC Patients. **A** Volcano plot of differentially expressed genes (DEGs) between bulk liver tissues from HCC and non-HCC patients. The red points indicate upregulated genes while the blue points represent downregulated genes (p.adj < 0.05 and |log_2_FC|> 1). The gray points represent genes with no significant difference. **B** DEGs between HCC tissues and peripheral blood mononuclear cells (PBMCs). The red points indicate upregulated genes while the blue points represent downregulated genes (p.adj < 0.05 and |log_2_FC|> 1). The gray points represent genes with no significant difference. **C** Overlap of DEGs across comparisons: HCC vs. Normal, Secreted proteins, CTC vs. HCC, and HCC vs. PBMC. **D** ROC (receiver operating characteristic) analysis of overlapped DEGs for distinguishing HCC from healthy individuals. **E** Coefficient profiles of shared DEGs derived from LASSO regression. **F** cross-validation curve for the penalty term. **G** ROC analysis of ADAMTS3 for distinguishing primary and recurrent HCC from healthy individuals. **H** Evaluation of ADAMTS3’s predictive performance in HCC patients with low AFP levels (bottom quartile) **I** ROC analysis of ADAMTS3 in an independent validation cohort. **J** Relative expression levels of ADAMTS3 in PBMCs, healthy individuals, individuals with obesity or non-alcoholic fatty liver (NAFL) disease, HCC patients, and circulating tumor cells (CTCs) from HCC patients. **K** Immune infiltration analysis comparing HCC samples with high (upper quartile) and low (lower quartile) ADAMTS3 expression. **L** Kaplan–Meier survival analysis of HCC patients with high versus low ADAMTS3 expression. ns indicates p > 0.05; * indicates p < 0.05; ** indicates p < 0.01; *** indicates p < 0.001.

### 3.2 ECM and immune remodeling in ADAMTS3-high HCC

To explore the mechanisms through which ADAMTS3 contributes to tumor progression and immune modulation, we conducted a comprehensive functional enrichment analysis comparing HCC bulk samples with high versus low ADAMTS3 expression (upper versus lower quartiles). Over-representation analysis indicated that elevated ADAMTS3 expression was strongly linked to extracellular alterations, particularly within the collagen-containing matrix— consistent with its established role in ECM regulation (Figure 3A). The top 10 enriched biological processes were predominantly matrix-related, including regulation of vascular endothelial growth factor signaling, elastic fiber assembly, and modulation of platelet activation (Figure 3B).

**Fig. 3.**
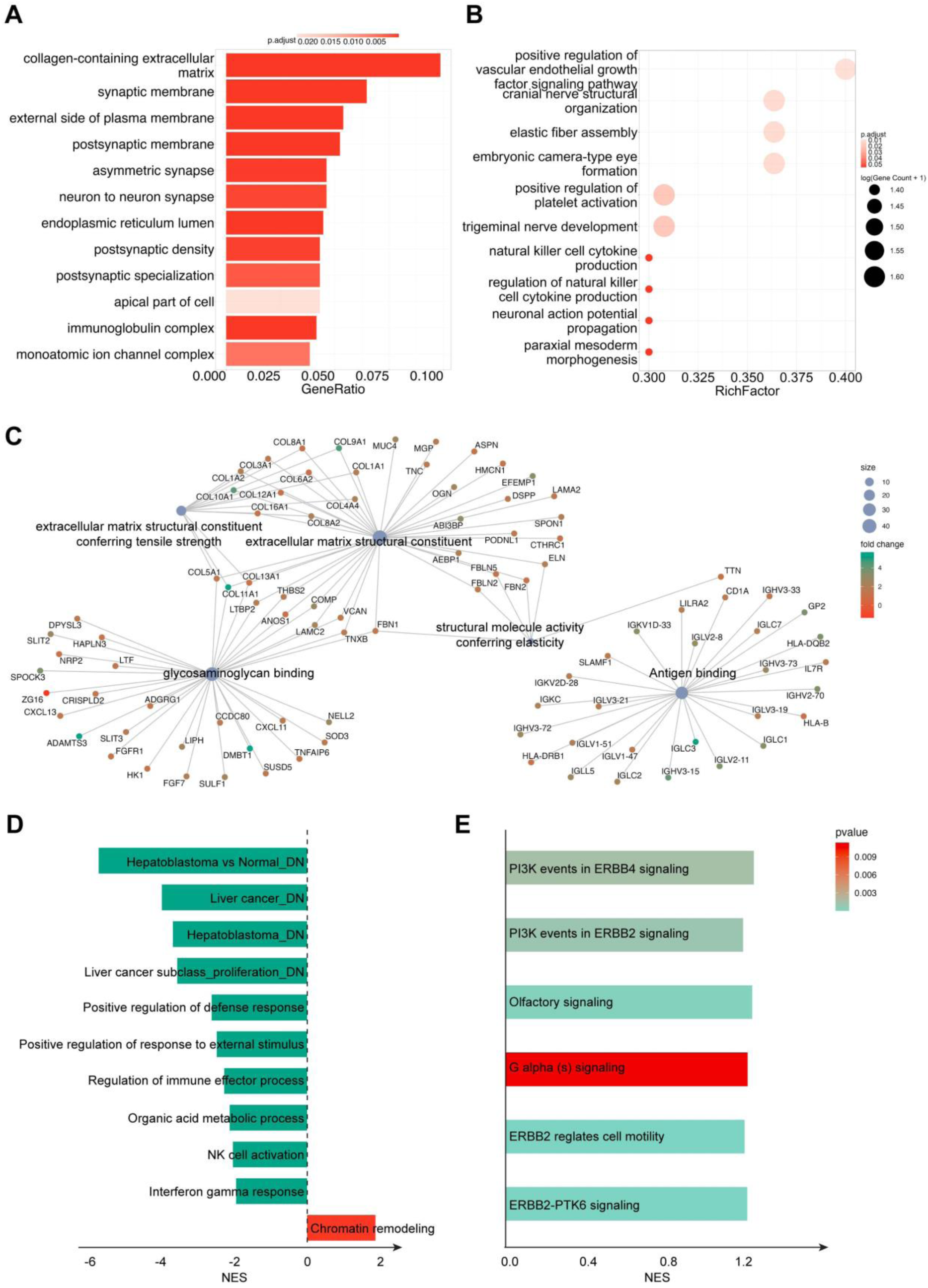
ECM and immune remodeling in ADAMTS3-high HCC **A** Overrepresented cellular components based on differentially expressed genes (DEGs) between ADAMTS3-high and ADAMTS3-low groups. **B** Overrepresented biological process based on DEGs between ADAMTS3-high and ADAMTS3-low groups. **C** Overrepresented molecular function in ADAMTS3-high versus ADAMTS3-low groups. **D** Top enriched features associated with ADAMTS3 high expression, as determined by GSEA. **E** Top enriched Reactome pathways of the DEGs.

In addition, enrichment of NK cell cytokine regulation suggests altered NK activity in ADAMTS3-high tumors. These immune changes appear to occur alongside ECM stiffness change and signaling alterations, as reflected by molecular function enrichment in structural molecule activity related to elasticity, ECM structural constituent, extracellular matrix structural constituent conferring tensile strength, and key signaling processes such as antigen binding (Figure 3C & Supporting Information - Figure S2A).

In line with these results, elevated ADAMTS3 expression was associated with macrophage polarization toward the immunosuppressive M2 subtype and diminished NK cell infiltration (Figure 2K), reflecting a reprogrammed immune microenvironment that may facilitate tumor progression. This was further supported by the marked suppression of defense responses, immune effector functions, NK cell activation, and interferon-gamma signaling observed in ADAMTS3-high HCC samples (Figure 3D). In addition, ADAMTS3-high HCC exhibits molecular signatures characteristic of highly proliferative HCC subsets, accompanied by elevated chromatin remodeling activity. To further explore these pathways, we performed heatmap analysis of the top 30 shared genes of these enriched pathways, highlighting distinct expression patterns associated with ADAMTS3 levels (Supporting Information - Figure S2B). To further elucidate the regulatory mechanisms through which ADAMTS3 drives proliferation and fosters an immunosuppressive tumor microenvironment, we examined the most significantly activated pathways in ADAMTS3-high samples. Strikingly, many of these pathways were membrane-associated, suggesting a role in transducing ECM stiffness alterations into intracellular biochemical signals (Figure 3E).

### 3.3 ADAMTS3 deficiency inhibits HCC Progression

To better delineate the role of ADAMTS3 in HCC progression, we generated ADAMTS3-knockdown Huh7 cell lines to assess its impact on cellular and biomechanical properties (Fig. 4A). ECM stiffness is a critical determinant of cancer cell behaviour—such as proliferation and metastasis potential—as well as key functions of surrounding immune cells, including activation, polarization, migration, infiltration, antigen presentation, and cytotoxicity^35–37^. ADAMTS3-induced changes in both intracellular and extracellular matrix stiffnesses. These are associated with immunosuppression and tumor progression. Therefore, the measurement of the cell stiffness can contribute to the general understanding of the role of ADAMTS2 in HCC. A reliable method to measure the cell stiffness relies on optical tweezers^38^ (see Methods). From our measurements, we observed that ADAMTS3 knockdown significantly reduced overall cellular stiffness, as reflected by a decrease in Young’s modulus. (Figure 4B-4D). Consistent with clinical association between high ADAMTS3 expression and a proliferative phenotype, knockdown of ADAMTS3 led to significantly reduced cell proliferation (Figure 4E). Furthermore, colony formation assays demonstrated impaired clonogenic capacity in ADAMTS3-deficient cells, indicating that ADAMTS3 contributes to both proliferative and clonogenic potential in HCC cells (Figure 4F). We next performed rescue experiments by restoring ADAMTS3 expression in knockdown cells to validate its role in driving HCC (Figure 4G). Restoration of ADAMTS3 expression effectively reversed the proliferation defects (Figure 4H) and partially rescued clonogenic ability (Figure 4I), further validating the role of ADAMTS3 in promoting HCC cell growth.

**Fig. 4.**
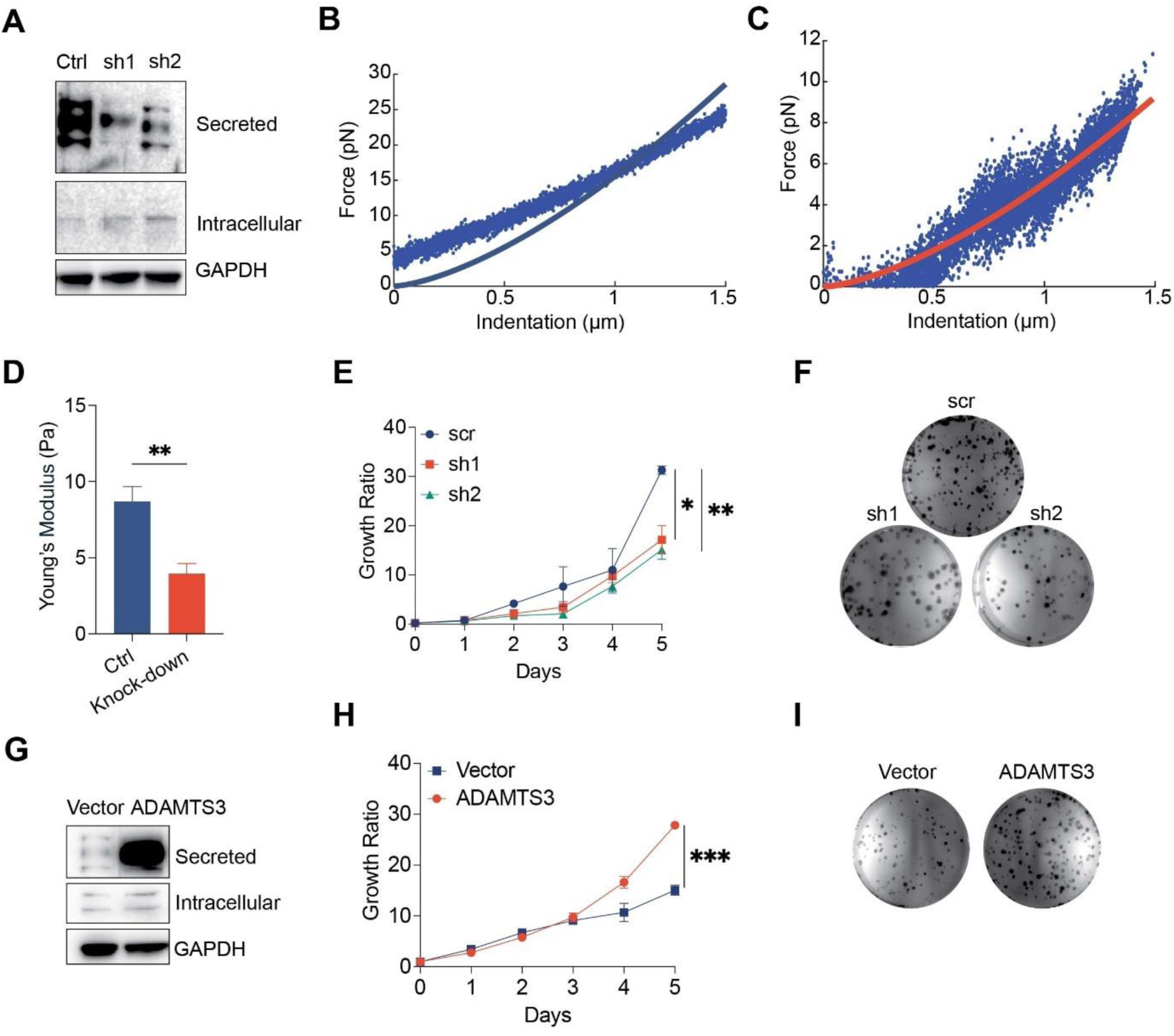
ADAMTS3 deficiency inhibits HCC Progression **A** Representative western blots of ADAMTS3 in wild-type or ADAMTS3 knock-down Huh-7 cells. **B** Force–indentation curve in wild-type Huh-7 cells measured by optical tweezers and fitted using the standard Hertz model. **C** Force–indentation curve in ADAMTS3 knock-down Huh-7 cells measured by optical tweezers and fitted using the standard Hertz model. **D** Quantification of Young’s modulus in wild-type and ADAMTS3 knockdown Huh7 cells. **E** Proliferation curves of wild-type and ADAMTS3 knockdown Huh7 cells; Cells were plated in normal growth medium in 96-well plates and CCK-8 assays were performed for a total of 5 days. **F** Colonies of wild-type and ADAMTS3 knockdown Huh7 cells as indicated. **G** Representative western blots of ADAMTS3 and GAPDH in ADAMTS3 knockdown Huh7 cells introduced either vector or ADAMTS3. **H** Proliferation curves of ADAMTS3 knockdown Huh7 cells introduced either vector or ADAMTS3; Cells were plated in normal growth medium in 96-well plates and CCK-8 assays were performed for a total of 5 days. **I** Colonies of ADAMTS3 knockdown Huh7 cells introduced either vector or ADAMTS3. Results presented are representatives from three independently conducted experiments. Growth curves were analyzed using two-way ANOVA followed by Tukey’s multiple comparisons test. Other statistical comparisons were performed using two-tailed Student’s t-tests.

### 3.4 ADAMTS3-GPCR Axis Fuels HCC

Motivated by these findings, we next aimed to investigate the functional role of ADAMTS3 in HCC progression by conducting transcriptome analysis on ADAMTS3-knockdown Huh7 cells and their corresponding scramble controls. Differential gene expression analysis revealed widespread transcriptional alterations following ADAMTS3 knock-down, with a distinct set of genes significantly up-or downregulated (Figure 5A; *p*.adj < 0.05, |Log_2_FoldChange| > 1). To gain insight into the functions and interrelationships of these transcriptional changes, we performed Gene Ontology (GO) over-representation analysis based on differentially expressed genes. Enrichment analysis of cellular components revealed prominent alterations in extracellular and membrane compartments, implicating ADAMTS3 in the structural organization of the cell–environment interface (Figure 5B). This is consistent with its known role as a regulatory checkpoint for collagen fibril assembly through enzymatic processing of procollagen^39^. We next examined the biological processes altered by ADAMTS3 deficiency and observed a significant enrichment of bone mineralization-related terms (Figure 5C), consistent with the well-established role of ADAMTS3 in skeletal development. Notably, enrichment of tissue remodeling and chemotaxis pathways points to additional functions of ADAMTS3 in shaping tissue structure and modulating cell motility. To further dissect the functional consequences of ADAMTS3 deficiency, we performed Gene Set Enrichment Analysis on molecular function level. Strikingly, the top-ranked molecular functions were uniformly downregulated in ADAMTS3-knockout cells (Figure 5D). These included multiple transcription-related activities, such as DNA-binding transcription factor activity, RNA polymerase II-specific transcription factor activity, sequence-specific double-stranded DNA binding, and cis-regulatory region binding. This overall reduction in transcriptional regulatory potential underscores a broader transcriptional suppression program following ADAMTS3 loss, which is consistent with the observation that HCC samples with high ADAMTS3 status is positively enriched of proliferative and chromatin remodeling pathways. These findings suggest that ADAMTS3 may indirectly support tumor growth by maintaining transcription factor activity necessary for cell cycle progression and survival. To further elucidate the downstream programs affected by ADAMTS3 loss, we performed hallmark gene set enrichment and pathway-based analyses. Gene set enrichment analysis revealed that ADAMTS3 deficiency significantly reversed aggressive phenotypes (Figure 5E). Notably, stem cell-associated downregulated genes were positively enriched (NES = 1.89, *p* < 0.001), suggesting a possible shift from a stem-like state to non-stem state. In line with this, metastasis-associated signatures were strongly suppressed (NES =-2.07, *p* < 0.001), along with downregulation of multicancer invasiveness (NES =-1.71, *p* < 0.05), indicating a reduction in invasive potential upon ADAMTS3 loss. Interestingly, apoptosis-related gene sets were upregulated (NES = 1.55, *p* < 0.01), consistent with decreased cell survival and proliferative capacity observed in earlier functional assays. Furthermore, the downregulation of the “assembly of collagen fibrils and other multimeric structures” signature (NES =-1.56, *p* < 0.05) underscores the structural disruption of the extracellular matrix following ADAMTS3 depletion. Complementary pathway enrichment analysis (Figure 5F) further confirmed these observations. Key signaling cascades were downregulated, including DAP12 signaling, GPCR signaling (including G alpha (s) signaling events), and nuclear receptor pathways, many of which are known to modulate immune and proliferative responses. Interestingly, GPCR is activated in HCC samples with high ADAMTS3 expression. Moreover, knock-down of ADAMTS3 in cell line reversed its activation, indicating the main role in transmitting ADAMTS3 signals to downstream nodes. In addition, pathways associated with RUNX2-mediated osteoblast differentiation and bone development were also suppressed, reinforcing the connection between ADAMTS3 and extracellular matrix homeostasis. The consistent downregulation of SUMOylation and collagen biosynthesis/modification pathways highlights a broader disturbance in post-translational regulatory mechanisms and matrix protein maturation.

**Fig. 5.**
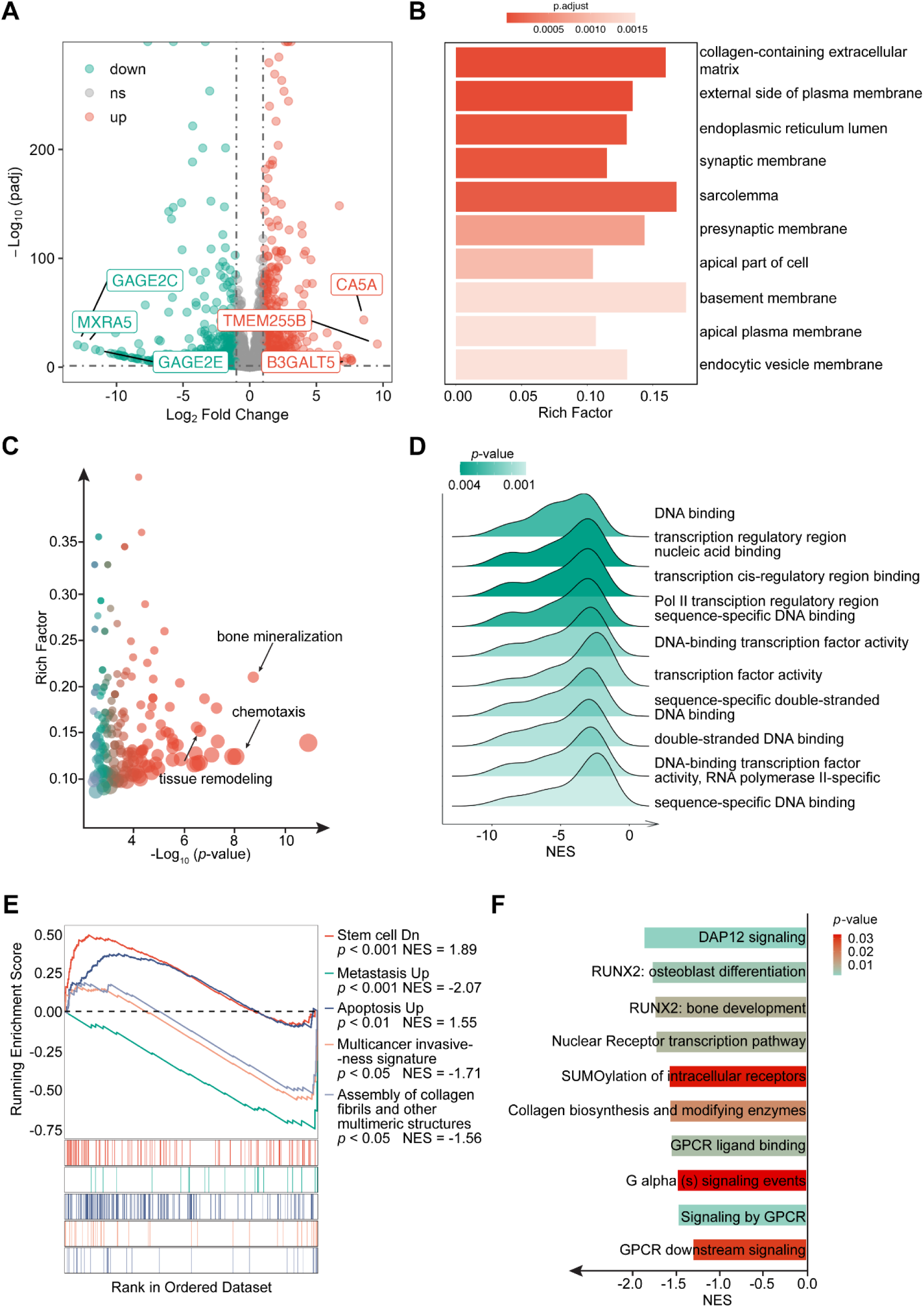
ADAMTS3-GPCR Axis Fuels HCC **A** Volcano plot of differentially expressed genes (DEGs) between ADAMTS3 knockdown Huh7 cells and its scramble control. The red points indicate upregulated genes while the green points represent downregulated genes (p.adj < 0.05 and |log2FC|> 1). The gray points represent genes with no significant difference. **B** Overrepresented cellular components based on DEGs due to ADAMTS3 knockdown. **C** Overrepresented biological processes based on DEGs due to ADAMTS3 knockdown. **D** Top enriched molecular features associated with ADAMTS3 deficiency, as determined by GSEA. **E** Top enriched phenotype features associated with ADAMTS3 deficiency, as determined by GSEA. **F** GSEA-based pathway enrichment analysis of ADAMTS3 deficiency

### 3.5 Detection of ADAMTS3 Using a DNA Origami-Based Biosensor

Serum biomarkers are typically present at very low concentrations, posing a significant challenge for traditional immunoassay-based detection methods. SERS offers a highly sensitive biosensing approach by leveraging the ability of metallic nanostructures to tightly confine electromagnetic fields, thereby enhancing Raman vibrational signals^40^. Therefore, after ADAMTS3 was identified as promising biomarker for HCC diagnosis, we fabricated a SERS probe and a SERS immunosensor for non-invasive detection of ADAMTS3 using a DNA Origami design comprising a dimer of Au nanoparticles (Au NPs) coupled to an aptamer specifically selected to identify the target biomarker. With respect to previously reported SERS biosensors for tumor biomarkers detection^41^, where classical antibodies were used, here we choose an aptamer as biosensing element. Aptamers are single-stranded synthetic DNA-or RNA-based oligonucleotides that fold into various shapes to bind to a specific target, which includes proteins, metals, and molecules. Aptamers have high affinity and high specificity that are comparable to that of antibodies and they can find important application in biomarker discovery^42–44^. In the design used here, two gold nanospheres are arranged as a dimer on a simple DNA origami scaffold to create strong SERS hotspots (Fig. 6A). An His-tag aptamer (see methods) is positioned in the gap between the nanospheres to enable specific binding of the target protein, ADAMTS3. One of the main advantages in using this specific design is the single detection spot, in principle able to achieve single-molecule detection^45^. With respect to SERS probes obtained from random distribution of metallic nanoparticles^41^, the DNA origami ensures a site-specific arrangement not only for the Au nanoparticles, but also for the aptamer in the gap where the SERS hot-spot is present. The only optimization needed in this case regards the choice of the proper molar ratio between the aptamer and the origami+NPs structure. We first characterized the structural morphology of this biosensor using transmission electron microscope (TEM). As shown in Figure 6B, the TEM images revealed the Au NPs arranged as a dimer with a narrow gap as expected by the design of the DNA origami template. Subsequently, the aptamer-functionalized DNA origami sensor was employed to detect purified recombinant His-ADAMTS3 protein. The detection process using the immunosensor requires only a few steps. First, it is important to note that the Au-NPs dimer generates a strong hot-spot in the gap region where the aptamer is located, even in the absence of the target biomarker. As a result, a background SERS spectrum corresponding to the presence of DNA (origami+aptamer) molecules is expected. Furthermore, in order to correctly assign the observed vibrational signals in the final biosensor, it is essential to characterize the Raman spectrum of the target protein ADAMTS3. For this purpose, we measured the Raman spectrum obtained from the protein deposited, with a concentration of 1 g/l, on a standard plasmonic substrate (prepared as a nanoporous Au layer^46^). The results, shown in Fig. 6C-I and 6C-II, highlight the distinction between the background spectrum (6C-I) and the ADAMTS3 spectrum (6C-II). Specifically, the background spectrum displays the characteristics DNA peaks at 1125, 1237, 1295, 1375 and 1453 cm^-1^ ^47^, whereas ADAMTS3 exhibits three distinct peaks at 1097, 1258 and 1328 cm^-1^ (Fig. 6C-II).

**Figure 6.**
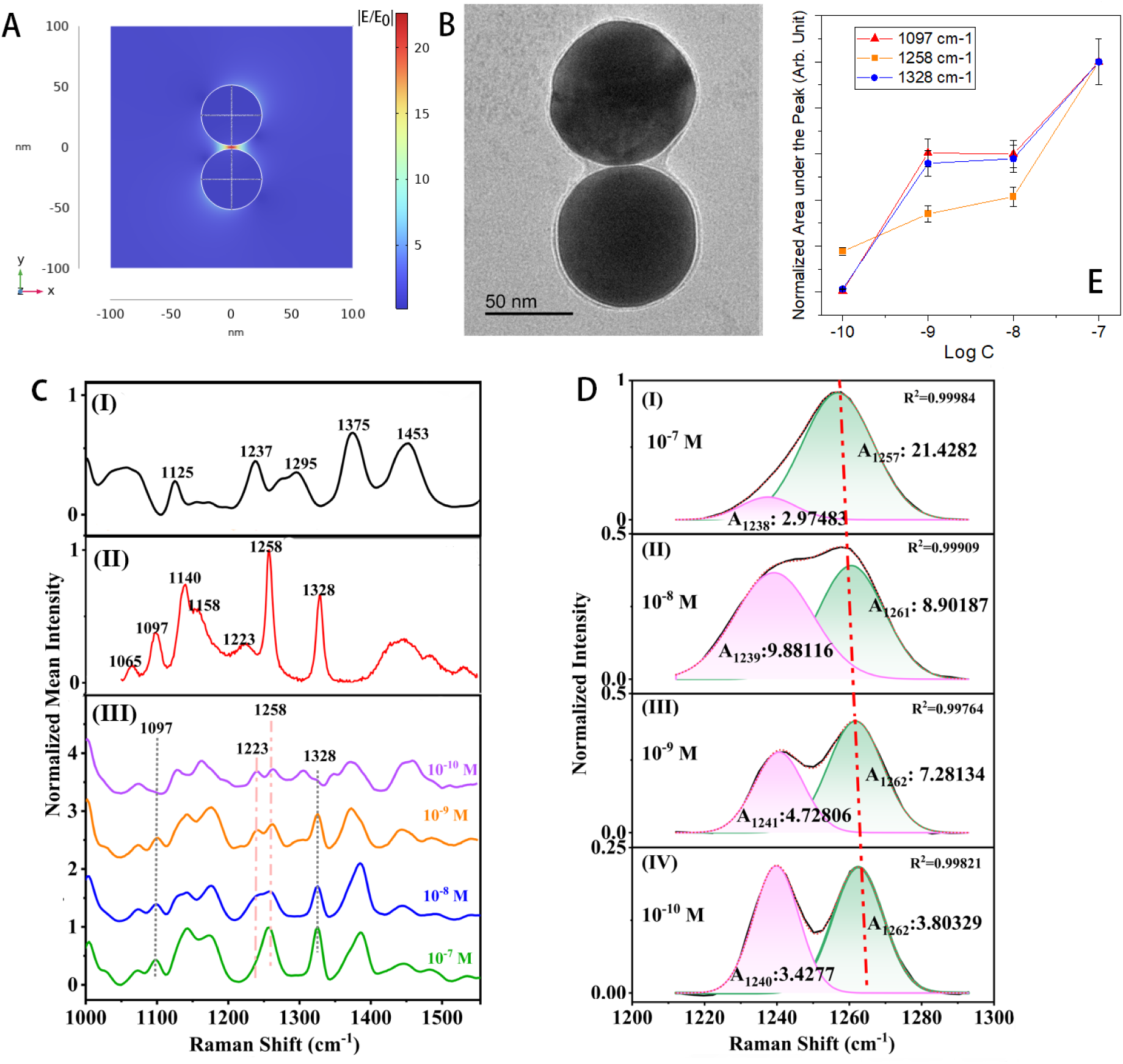
A. Numerical simulation illustrating the electromagnetic field confinement / enhancement between the Au NPs. **B.** TEM micrograph of two Au NPs self-assembled on the origami structure. **C.** blank reference of the DNA origami+aptamer platform **(I)**; high-intensity signal of the ADAMTS3 deposited on a nanoporous Au film with a concentration of 1g/l **(II);** Comparison of SERS signals across substrates: concentration-dependent series on DNA origami **(III)**. **D.** Example of deconvoluted peaks of the normalized SERS spectra (1210-1295 cm⁻¹), labeled with their relative areas. Notably, the peak fitted with the red dotted line shifts with varying concentration. **E.** Scatter plot of the ‘area under the peak vs. concentration’ for the three main characteristics peaks of ADAMTS3.

We next carried out biosensing experiments by incubating the origami sensor with ADAMTS3 protein at varying concentrations. As shown in Fig. 6C-III, the sensor demonstrated detection capability down to 10⁻¹⁰ M. To properly analyze the spectra, peak deconvolution was performed to account for overlapping DNA signals (Fig. 6D). This allowed us to generate scatter plots of peak intensity versus concentration for the three characteristic ADAMTS3 peaks (Fig. 6E). A clear linear response was observed across all evaluated wavenumbers, with the strongest sensitivity achieved at 1258 cm^-1^.

## 4. Conclusions

In summary, this study identifies ADAMTS3 as a novel serum biomarker for the early diagnosis of HCC. Our findings reveal a previously unrecognized role for ADAMTS3 in promoting HCC progression and mediating immune suppression via ECM stiffness-regulated signalling pathways.

By screening for upregulated secreted proteins in a discovery cohort comprising non-HCC individuals—including healthy controls and patients with obesity or NAFLD—and HCC patients with diverse etiologies (HBV, HCV, and non-viral) and disease stages, we identified several biomarker candidates including ADAMTS3. Further evaluation of its performance in distinguishing HCC demonstrated its strong potential for early diagnosis, with an AUC of 0.9039 (95% CI: 81.79%–99.00%) for primary HCC and 0.8704 (95% CI: 77.32%–96.75%) for recurrent HCC. In contrast, the most prevalent serum protein marker used in clinical practice, AFP, shows an AUC ranging from 0.75 to 0.82 ^48^, and other two approved biomarkers, DCP and AFP-L3, had the AUCs of 0.72 and 0.66, respectively ^49^. In addition, ADAMTS3 also exhibits an AUC of 0.8765 (95% CI: 78.36%-96.94%) for AFP-negative HCC, outperforming currently available HCC biomarkers and holds strong promise to improving HCC early diagnosis.

In physiological states, ADAMTS3 mediates proteolytic processing of procollagen II and fibronectin, which is essential for fibril formation and ECM remodeling ^39,50–52^. Beyond this structural role, ADAMTS3 also directly regulates the activity of ECM-associated receptor by cleavage their ligands, thereby promotes embryonic lymphangiogenesis and regulates placental angiogenesis ^16,50,53^. In cancerous contexts, although limited research has implicated multifaceted roles for ADAMTS3–including tumor-promoting behavior in glioma stem cells and tumor-suppressive effect in breast cancer–its role in HCC remains poorly understood and has not been systematically investigated ^51,54^. Our study fills this critical gap by demonstrating that ADAMTS3 is not only a promising biomarker for early HCC diagnosis but also an active driver of tumor progression. Through comprehensive analysis of transcriptomic profiles of HCC patients stratified by ADAMTS3 expression levels, we revealed that ADAMTS3 overexpression correlates with the ECM remodeling and increased stiffness of both the ECM and cellular components, which promotes immunosuppression and likely activates downstream mechano-transduction pathways that support cancer cell proliferation ^55^. Further knockdown of ADAMTS3 in HCC cell lines resulted in decreased cellular stiffness, along with impaired proliferation and colony-forming ability. Transcriptomic analysis following ADAMTS3 knockdown revealed downregulation of collagen fibril assembly and other multimeric structural components, along with the loss of stemness and metastasis-associated gene signatures, consistent with the impaired stiffness and proliferative phenotype observed in ADAMTS3-deficient HCC cell lines. Moreover, signaling cascades related to G protein–coupled receptor (GPCR) activity, particularly Gαs-mediated pathways, were markedly suppressed upon ADAMTS3 knockdown. These findings highlight a mechanistic link between ADAMTS3-mediated ECM remodeling and downstream GPCR signaling, which may underlie the observed changes in tumor cell behavior and immune microenvironment modulation.

Therefore, this study expands the functional landscape of ADAMTS3, positioning it as a molecular integrator that couples ECM structural dynamics with cellular signaling and immune regulation in HCC. Promising discriminating power and mechanistic driver role of oncogenesis makes ADAMTS3 an ideal biomarker for HCC diagnosis.

Finally, here we also proposed a DNA origami–based biosensor capable of high-sensitive and specific detection of ADAMTS3. By engineering stable and reproducible SERS-active plasmonic nanostructures through DNA origami self-assembly, this platform achieves picomolar-level sensitivity, paving the way for early-stage HCC screening in at-risk populations.

## Acknowledgements

The authors acknowledge financial support from National Key Research and Development Project of China, Grant No. 2023YFF0613603 and National Natural Science Foundation of China, Grant No. 22202167.

## Contributions

Conceptualization and design of the study: C. Wang and S. Jin. Acquisition of data: C. Wang, H. Jin, Q. Zhang, Y. Zou. Methodology: C. Wang, Q. Zhang, Y. Wang. Modelling: A. Douaki and N. Maccaferri. Analysis and interpretation of data: C. Wang, Y. Wang, Z. Fang, L. Yao. Y. Cai, H. Jin. Writing – original draft & submitted manuscript: C. Wang. J. Feng, D. Garoli. Study supervision and funding acquisition: H. Jin, S. Jin

